# A dataset containing speciation rates, uncertainty, and presence-absence matrices for more than 34,000 vertebrate species

**DOI:** 10.1101/2024.04.09.588748

**Authors:** Juan D. Vásquez-Restrepo

**Affiliations:** Laboratorio de Herpetología, Museo de Zoología “Alfonso L. Herrera”, Universidad Nacional Autónoma de México, Ciudad de México, México; Department of Ecology and Evolutionary Biology, Princeton University, Princeton, United States

## Abstract

The increasing availability of phylogenetic data has facilitated the exploration of macroecological and macroevolutionary patterns across diverse spatial and temporal scales. However, calculating some model-based diversification metrics often requires significant computational power and time. To address this, I present a comprehensive dataset of Bayesian diversification rates for over 34,000 vertebrate species, spanning five major groups: amphibians, birds, mammals, reptiles, and sharks. This is the first large-scale dataset of its kind, providing both continuous and binary data for speciation rates and presence-absence matrices, respectively, with global coverage. The dataset not only enables analyses of evolutionary and spatial diversity patterns but also democratizes access to data-intensive studies. Additionally, as it is based on Markov chains, the dataset can be customized and extended without the need to start from scratch, offering flexibility for future research on diversification dynamics.

## Introduction

Understanding how diversity vary in time and space is one of the most important questions in ecology and evolutionary biology, as delving into the causes and consequences of disparity can offer valuable insights into the origin and the future of species^1^. Far from having a single or simple answer, several hypotheses have been proposed to explain why some lineages or geographic regions are more prolific or prone to accumulate greater diversity than others. These include factors such as evolutionary time, area, spatial heterogeneity, climate stability, mutation rates, and competition^2,3,4,5^. However, exploring large-scale patterns is a data-intensive task that, in the case of diversification dynamics, requires both phylogenetic trees and robust mathematical frameworks to infer rates of shift^6^.

Previous studies have generated “fully sampled” phylogenies or supertrees for the major vertebrate radiations^7,8,9,10,11^ and used them to calculate diversification rates, most of them, through deterministic approaches. Although computationally fast, these methods do not account for variable rates within the tree and are limited to speciation-only. In contrast, probabilistic approaches (model-based) enable the inference of more complex evolutionary scenarios^1^.

However, applying such methods to supertrees significantly increases the computational burden and requires additional programming skills. Moreover, some methods are sensitive to taxon sampling, limiting them to comprehensive species-level phylogenies rather than higher-rank trees^12^. For this reason, studies using probabilistic methods are often restricted to smaller taxonomic subsets. In addition, another issue arising from supertrees is their high level of unresolved groups^13^, which can affect the final estimates if phylogenetic uncertainty is not taken into account^14^. This is particularly critical for inferring the evolution of traits or for biogeography, as randomly resolved polytomies disrupt natural patterns, but in the absence of real data, they seem remain useful for inferring evolutionary rates^15^.

In this study, I explore the space of possibilities using a Bayesian approach for estimating diversification rates in the five major groups of vertebrates. This dataset is the result of an exploratory analysis driven by a genuine curiosity, but for which, I currently have no further plans for its use in the short term. For this reason, and considering that the results meet a minimum performance threshold to be deemed reliable, I would like to make it publicly available. My major motivation is not to be the “dog in the manger”, a metaphor describing a person who has no need for, or ability to use a possession that would be of use or value to others, but prevents them from having it. This dataset could be a valuable resource for researchers interested in macroecology and macroevolution, whether they choose to use it as it is or as a foundation for further analyses, ultimately, saving time and effort. Here, I provide a brief description of the data but will not delve into the causes or consequences of the observed biodiversity patterns in detail.

## Methods

### Phylogenetic sampling

I used available “fully-sampled” time-calibrated supertrees for five monophyletic groups of extant vertebrates: amphibians^7^ (subclass Lissamphibia: 7,238 species), birds^8^ (class Aves: 9,993 species), mammals^9^ (class Mammalia: 5,911 species), reptiles^10^ (order Squamata: 9,754 species), and sharks^11^ (class Chondrichthyes: 1,190 species). Since these phylogenies were inferred using a mixed approach that combines a base topology derived from species with genetic data and the imputation of species lacking such information, the resulting consensus tree is highly polytomic. To address this issue, I subsampled 100 randomly dichotomized trees for each group and performed all analyses on this set of replicates^14^. The trees were obtained from the VertLife website (http://vertlife.org).

### Diversification rates

Although this dataset primarily focuses on probabilistic diversification rates, I also calculated them using a commonly and widely used deterministic approach to compare their variability and provide two different alternatives, as described below:

#### BAMM

The Bayesian Analysis of Macroevolutionary Mixtures (BAMM) uses Markov Chain Monte Carlo (MCMC) to model complex dynamics of speciation (λ) and extinction (µ), allowing for heterogeneity in evolutionary rates^16^. For each of the 100 trees in the five vertebrate groups, I ran a BAMM analysis of 2 × 10^7^ generations, with four MCMC chains and sampling every 1000 iterations for a total of 20,000 samples, using ‘BAMM’^16^ v.2.5.0. Default control file was implemented with custom priors obtained for each tree using the function *setBAMMpriors* in R’s package ‘BAMMtools’^17^ v.2.1.10. Due to the computationally demanding nature of this analysis, it was run on a server equipped with 40 Intel Xeon Gold 6230 2.10 GHz processors and 125 GB of RAM. Additionally, to optimize resources, each set of 100 runs was partitioned into four parallel processes of 25 replicates. The BAMM outputs were analyzed and processed locally. A script for running multiple BAMM analyses automatically can also be found at GitHub (https://github.com/VR-Daniel/Multiple-BAMM).

#### DivRate

The DivRate index aims to capture clade-wide “diversification” estimates based on the branching patterns of birth-only and homogeneous birth-death trees^8,18^. Despite its name, DivRate does not represent a net diversification estimate (speciation + extinction) but instead focuses solely on speciation^19^. DivRate is calculated as the inverse of the Evolutionary Distinctiveness (ED), which measures the evolutionary isolation of a species^19,20^,21. The ED metric can be calculated using two approaches: the “fair proportion”, which divides branch lengths proportionally among all descendant species, or the “equal splits”, which divides branch lengths equally among all descendant clades at each branching point ^22^. To compute the ED, I used the *evol_distinct* function in the ‘phyloregion’^23^ package, which is computationally more efficient than other libraries.

### Matrices of presence-absence

To facilitate data spatialization (Fig. 1), I generated presence-absence matrices for each vertebrate group using distribution polygons from the International Union for Conservation of Nature (IUCN, https://www.iucnredlist.org) as in the 2023-1 update: amphibians (7,958 species), mammals (5,884 species), reptiles (9,821 species), and sharks (1,201 species). For birds (10,920 species), I employed a previously acquired dataset from an earlier update (2021-3) due to restricted data access. First, I corrected invalid geometries in the polygons using the built-in algorithm in QGIS v.3.28 for fixing geometries. This algorithm employs the structure method, which aims to repair polygon’s topology while preserving node integrity. Next, I generated the presence-absence matrices using the *lets*.*presab* function in ‘letsR’^24^ v.5.0 at a spatial resolution of one degree in geographic coordinates (WGS84). To spatialize the speciation rates accounting for species richness, I standardized the rates as the ratio between the mean diversification rates per cell and the weighted mean across all cells, using species richness as the weighting variable. Values above one indicate speciation rates higher than the average, while values below, indicate rates lower than the mean.

**Figure 1.**
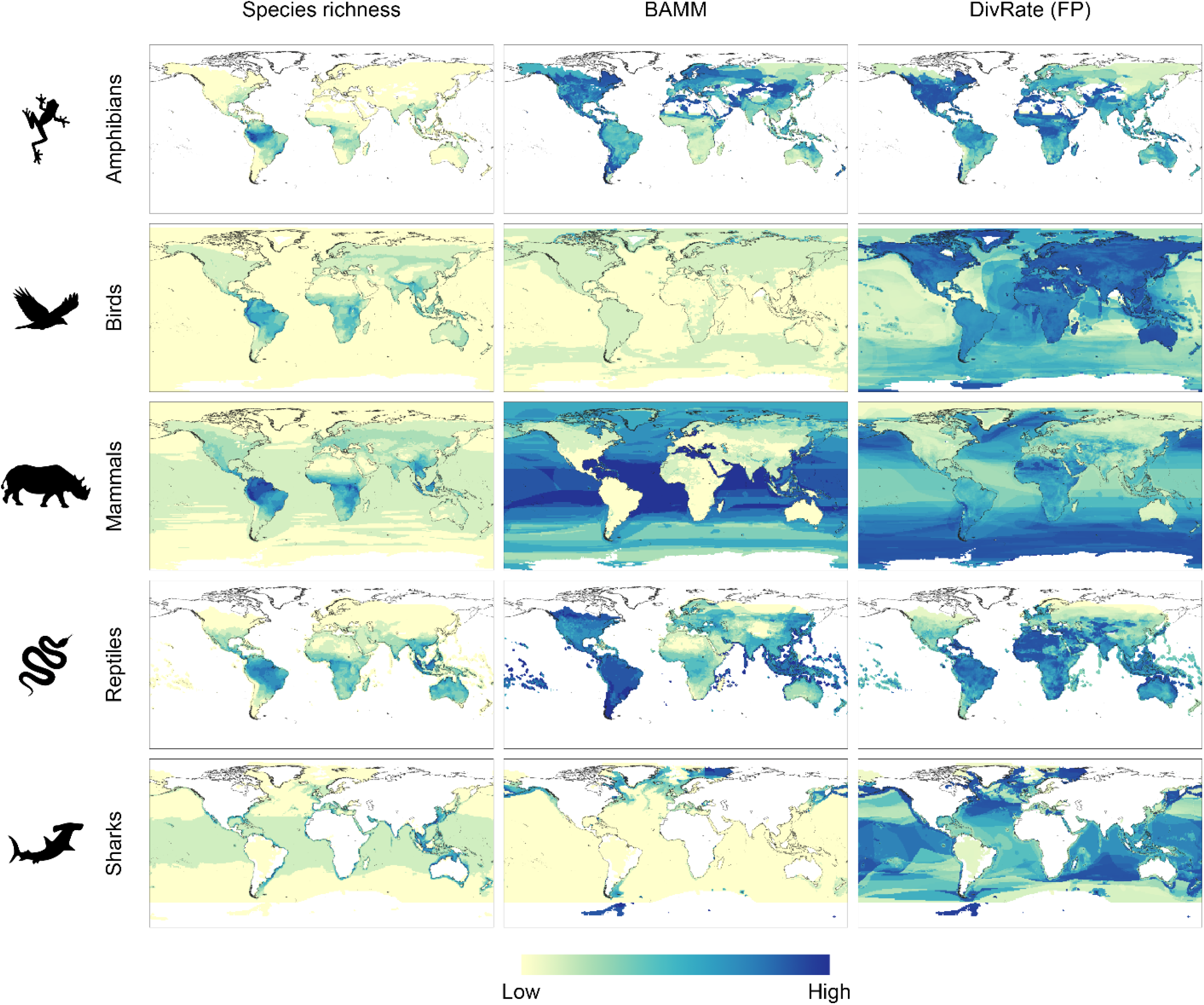
Geographic spatialization of species richness and speciation rates, based on the generated presence-absence matrices and diversification metrics. Maps are quantile-based colored.

### Taxonomic harmonization

Because the binomial names of species may vary across time and datasets, I included the GBIF usage key as a unique numeric identifier for each taxon. GBIF keys are invariant IDs associated with specific taxonomic ranks, enabling species tracking even when names become invalid. To assign these keys, I used GBIF’s built-in Look Up Tool to match species names from phylogenetic trees and IUCN polygons with GBIF’s backbone taxonomy.

This tool operates on three levels: (1) EXACT matches, where the provided name and GBIF’s name are identical; (2) FUZZY matches, when the names differ slightly (e.g., due to grammatical gender discordance or grammar mistakes); and (3) HIGHERRANK matches, when no direct match is found. For the diversification rates dataset, no matches with GBIF’s backbone taxonomy included 11 amphibians (0.15%), 6 reptiles (0.06%), 2 birds (0.02%), 9 mammals (0.15%), and 6 sharks (0.5%). Similarly, for the presence-absence matrices, no matches included 26 amphibians (0.33%), 4 reptiles (0.04%), 1 bird (0.01%), 2 mammals (0.03%), and 2 sharks (0.17%). To address non-matching species between datasets, I applied an additional string similarity test to reduce errors caused by misspelled names or ambiguities in taxonomy. This procedure uses the Optimal String Alignment algorithm based on the Levenshtein distance, to calculate the number of deletions, insertions, and substitutions required to convert one string into another while allowing adjacent character transpositions. For this, I built a custom script which is available on GitHub (https://github.com/VR-Daniel/taxTools).

### Data records

A summary list of the files included in this dataset, their description, format, and content is presented in Table 1. The data can be accessed and downloaded from Figshare: http://doi.org/10.6084/m9.figshare.24894363.

**Table 1.** Files’ structure, names, formats, and content of the elements composing this dataset.

| File | Format | Description |
| --- | --- | --- |
| **1. Diversification** | **.zip** | **Main folder** |
| GROUP |  | Folder |
| Branch lengths |  | Subfolder |
| *GROUP_BRANCH LENGTHS* | .xlsx | Branch lengths file for the 100 trees and summary statistics by group (in Species/Myr) |
| Speciation rates |  | Subfolder |
| *GROUP_BAMM* | .xlsx | BAMM estimates file for the 100 trees and summary statistics by group (in Species/Myr) |
| *GROUP_DivRate_ES* | .xlsx | DivRate “Equal Splits” estimates file for the 100 trees and summary statistics by group (in Species/Myr) |
| *GROUP_DivRate_FP* | .xlsx | DivRate “Fair Proportion” estimates file for the 100 trees and summary statistics by group (in Species/Myr) |
| *GROUP_BAMM FILES* | .zip | Raw BAMM outputs, including the runs info, mcmc outputs, event data, chain swap, and the 100 species trees |
| *GROUP_PRIORS* | .RData | Prior list for each run/tree (must be loaded using *readRDS*) |
| *GROUP_SAMPLED TREES* | .nex | Species trees |
| *divcontrol_TEMPLATE* | .txt | Control file template for the BAMM analyses. |
| **2. Presence-Absence** | **.zip** | **Main folder** |
| *paMa_GROUP* | .RData | Presence-Absence matrices by GROUP (can be loaded using *load*) |
| **3. Rasters** | **.zip** | **Main folder** |
| Richness |  | Subfolder |
| *Richness_GROUP* | .tiff | Species richness per GROUP (at 1°) |
| Speciation rates |  | Subfolder |
| *GROUP_BAMM* | .tiff | Raster file for the standardized BAMM speciation rates (at 1°) |
| *GROUP_DivRate_FP* | .tiff | Raster file for the standardized DivRate “Fair Proportion” speciation rates (at 1°) |
| **4. Dictionaries** | **.zip** | **Main folder** |
| paMa_Taxonomy | .xlsx | Harmonized taxonomy for the presence-absence dataset |
| phylo_Taxonomy | .xlsx | Harmonized taxonomy for the diversification rates dataset |

### Technical validation

To validate the MCMC convergence, I performed the Heidelberger and Welch’s test^25^, which assess stationarity in Markov chains by iteratively splitting them and testing if the means of consecutive segments are stable. For this purpose, I used the function *heidel*.*diag* in the ‘coda’^26^ library v.0.19-4.1 with the default parameters. Moreover, because trees were randomly dichotomized, they differ in topology and are therefore unlikely to converge to the same likelihood value. Nevertheless, if each tree converges independently to its most optimal solution, a mean convergence pattern rather than random variation is expected. This was visually assessed through a mean log-likelihood trace plot, using the replicates that passed the Heidelberger and Welch’s test (hereafter passed runs) per group and discarding the first half of the iterations as burn-in (Fig. 2). Subsequently, I calculated the Effective Sample Size (ESS) for the subset of passed runs, both for the number of shifts and log-likelihood parameters, using the *effectiveSize* function in the same *op. cit*. library (Fig. 2). A major challenge when analyzing large datasets is that, as tree size increases, the number of generations required to adequately sample the posterior distribution also increases, significantly but unpredictably. Consequently, increasing the number of generations *n-times* through trial and error would be computationally unfeasible in terms of cost-benefit. However, because ESS evaluates sample independence, running multiple independent BAMM analyses can help explore different peaks in the parameter landscape by generating a distribution frequency, compensating for variation in ESS among runs. In this case, I used a permissive minimum threshold (ESS ≥ 100). Based on the above, all the results and summaries herein presented correspond to the number of replicates with sufficient performance per group.

**Figure 2.**
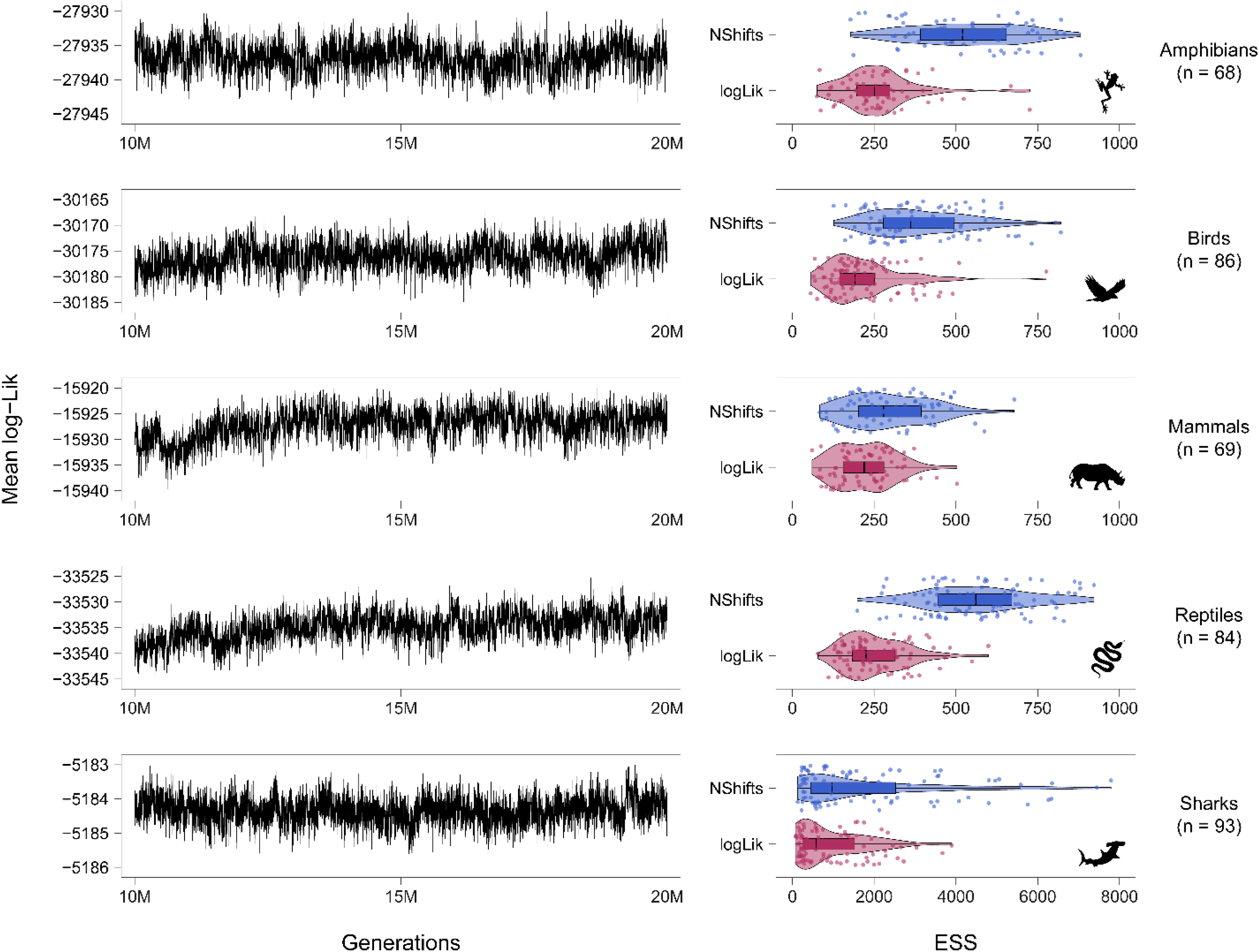
Mean log-likelihood trace plots and ESS values distribution for the five vertebrate groups included in this dataset. The numbers in parenthesis indicate the number of retained passed runs with estimated parameters’ ESS equal or greater than 100.

### Usage notes

Far from being perfect or the only option^1,27^, the method I chose to employ may reflect a personal preference, since it has been demonstrated to be robust and consistent, staying afloat despite the critics^12,28,29,30^. It also has extensive and detailed documentation, making it easy to implement, and benefits from an active community willing to assist with troubleshooting. However, users must be aware that diversification rates are known to be method-dependent^27^, which is why it is recommended that they first familiarize themselves with the calculation and interpretation of diversification metrics before using them.

In many analyses, diversification rates often lack measures of uncertainty. In this dataset, the inclusion of replicates helps, in part, to address this issue. Particularly, BAMM seems to be more influenced by the random dichotomization of the trees than DivRate (Fig. 3a–c). However, except for sharks, most of the variation for the other groups remains below the 30% threshold. Herein, I focused on speciation rather than extinction rates, as extinction rates based solely on phylogenetic data from living species can be biased, despite they may be correlated with true rates^16^. However, since I included all BAMM output files, users interested in additional information, such as extinction rates or the *a posteriori* maximum probability shift configuration for each run, can easily extract it using the *getEventData* function in the ‘BAMMtools’ package (refer to the package’s help files for further details). Users can also modify the parameters of each run or increase the number of generations without rerunning the entire analysis from scratch (previous experience with BAMM input files is required). To do so, they must refer to the ‘divcontrol_TEMPLATE’ file and set the argument *loadEventData = 1*.

**Figure 3.**
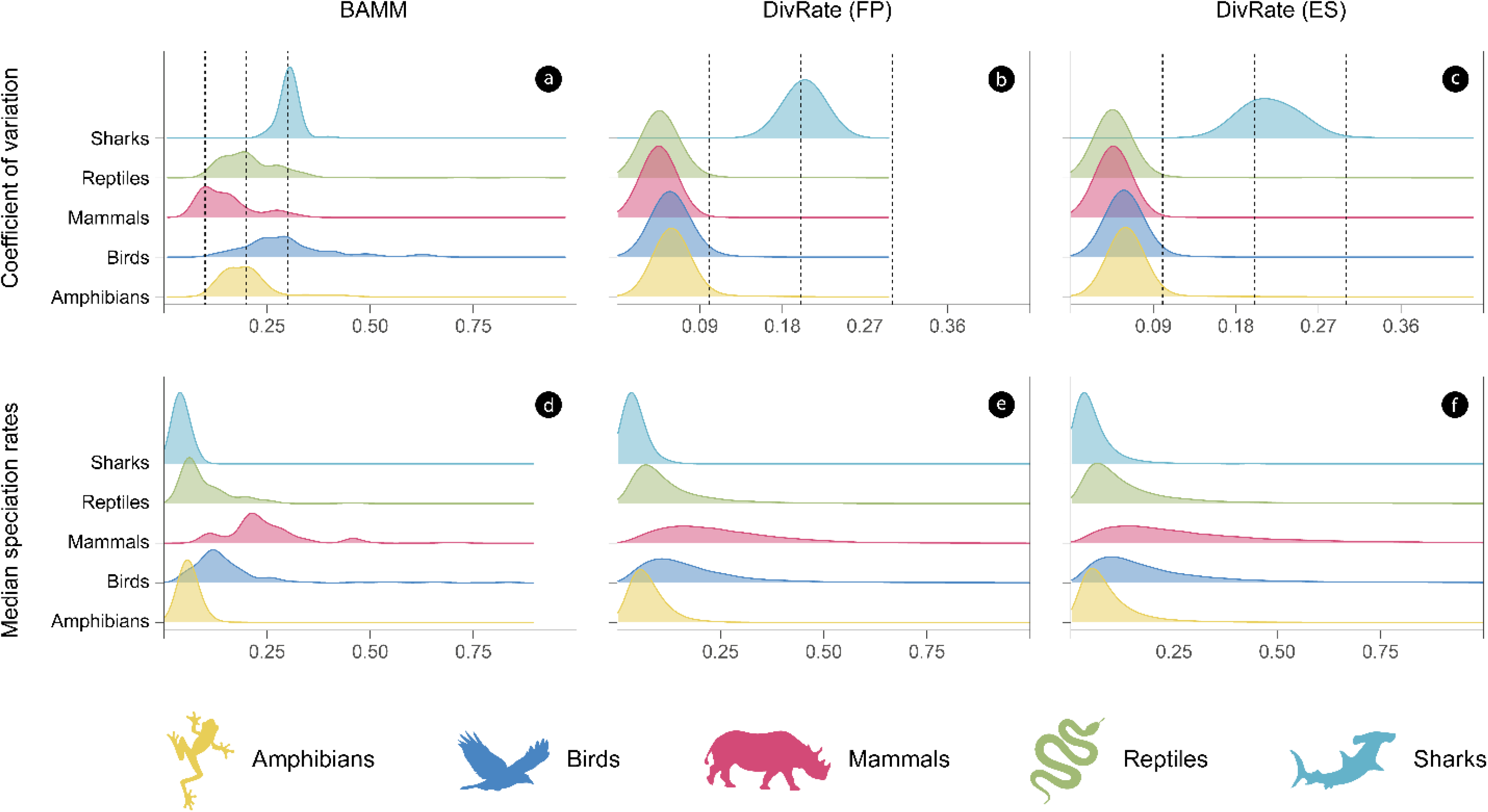
Distribution of the variation in speciation rates (a–c) and median speciation rates (d–f) for the five groups of vertebrates included in this work. The vertical dashed lines indicate the 10%, 20%, and 30% thresholds, respectively.

On the other hand, taxonomic discrepancies between diversification metrics and presence-absence matrices may arise due to differences in publication dates and data sources. To ensure accurate data matching, users should harmonize the taxonomy of their species of interest. The provided GBIF usage keys, which are consistent across datasets, offer a simple and effective solution for most cases. However, as these IDs were matched using a heuristic method, some mismatches may still occur. When spatializing data, users should also consider the right-skewed nature of diversification metrics (Fig. 3d–f) and the variation in species richness per pixel, as these can affect central tendency measures such as the mean. Alternatives like the median or geometric mean, along with standardized relative measures, are often more appropriate. Careful consideration of these factors will ensure more robust and meaningful inferences.

## Code availability

All the code and software used was fully described in the Methods section.

## Acknowledgements

I would like to thank Julián A. Velasco and the *Instituto de Ciencias de la Atmósfera y Cambio Climático* at *Universidad Nacional Autónoma de México*, for allowing me to use their informatic infrastructure, which was acquired supported by the DGAPA grant UNAM-PAPIIT IA201320.

